# Crystal structures of Scone, pseudosymmetric folding of a symmetric designer protein

**DOI:** 10.1101/2021.04.12.439409

**Authors:** Bram Mylemans, Theo Killian, Laurens Vandebroek, Luc Van Meervelt, Jeremy R.H. Tame, Tatjana N. Parac-Vogt, Arnout R.D. Voet

## Abstract

Recent years have seen a raise in the development of computational proteins including symmetric ones. We recently developed a nine-fold symmetric *β*-propeller protein named Cake. Here we wanted to further engineer this protein to a three-fold symmetric nine-bladed propeller using computational design. Two nine-bladed propeller proteins were designed, named Scone-E and Scone-R. Crystallography however revealed the structure of both designs to adopt an eight-fold conformation with distorted termini, leading to a pseudo-symmetric protein. One of the proteins could only be crystallized upon addition of a polyoxometalate highlighting the usefulness of these molecules as a crystallisation additive.

## 2 Introduction

Protein design has come a long way since its emergence in the late 80s when the first artificial proteins, consisting of helical bundles, were created by the group of De Grado Ho and DeGrado [1987], Korendovych and DeGrado [2020]. Amino acid selection at that time was guided purely by physicochemical principles, since the database of known protein structures at that time was still comparatively small. As the understanding of protein folding improved, new computational techniques (such as side-chain repacking) were developed that refined the modelling and design procedure. These algorithms automated the process Dahiyat and Mayo [1996], allowing a small natural protein domain to be repacked without human input Dahiyat and Mayo [1997]. As many more protein structures became available through the Protein Data Bank, fragment-based methods could be developed that eventually led to the first protein with a novel topology, called TOP7 Kuhlman et al. [2003]. Structure databases continued to expand as computers and synthetic gene synthesis became faster and cheaper, allowing the field to grow steadily. With increased efforts to relate protein sequences to structure in a predictive fashion, and the creation of freely available software such as the Rosetta suite Leaver-Fay et al. [2011], protein design evolved from a purely academic field to application-driven development. Recent successes include the creation of protein logic gates for gene regulation, the design of a vaccine for Respiratory syncytial virus and a protein inhibitor of SARS-CoV-2 Chen et al. [2020], Sesterhenn et al. [2020], Cao et al. [2020].

Many different globular protein folds have been targeted for (re)design, but monomeric folds with internal symmetry have been especially popular, including the *β*-trefoil fold Lee and Blaber [2011], TIM-barrels Huang et al. [2016] and *β*-barrels Dou et al. [2018]. The use of symmetry reduces the sequence search space during the design step as the protein can be assembled from identical repeats. The symmetry can also assist in the bottom-up design of more complex structures such as capsids, arrays or frameworks Hsia et al. [2016], Gonen et al. [2015].

Another protein fold with internal symmetry is the *β*-propeller fold, which consists of small *β*-sheets (each with four strands) arranged around a central channel. Natural propeller proteins contain between four and ten repeats of this motif, also called a blade Fülöp and Jones [1999]. The first successful design of a *β*-propeller was the six-bladed Pizza protein Voet et al. [2014] which has six identical repeats. Interestingly, this protein could self-assemble from two- or three-bladed fragments, as a trimer or dimer respectively. The Pizza protein was further functionalized to biomineralize a cadmium chloride nanocrystal Voet et al. [2015], to form multimeric assemblies by incorporation of coiled-coils Vrancken et al. [2020], to bind inorganic polyoxometalates (POMs) that control the crystal packing Vandebroek et al. [2020], and to act as a novel hydrolase Clarke et al. [2019]. These examples show the potential of the *β*-propeller fold as a functional building block for a variety of materials. Other symmetric proteins have been designed, such as the eight-fold symmetric Tako8 protein and its four-fold variant Ika8. The Tako8 protein is unable to assemble from smaller repeat fragments because of repulsion between the charged blades. To overcome this effect, the protein was redesigned with alternating repeats, compensating the charges. This restored the ability to self-assemble and improved overall stability Noguchi et al. [2019]. Most recently we designed the Cake protein, which can adopt eight-fold or nine-fold symmetry depending on the number of repeats expressed. While this structural plasticity has shed light on the evolutionary mechanisms, yielding diverse repeat numbers in natural *β*-propellers, it may lead to unexpected results during further functionalization of the Cake protein as a building block Mylemans et al. [2020a]. We therefore set out to redesign Cake9 into a three-fold symmetric nine-bladed protein.

## 3 Materials and Methods

### 3.1 Computational Protein Design

The design started from the protein with PDB entry 3hxj, the same protein that was used in the design of the Cake protein Mylemans et al. [2020a]. A three repeat fragment was isolated. From this fragment a three-fold symmetric template was created using the SymDock protocol from Rosetta André et al. [2007]. The order of the repeats four to seven was permuted in all possible ways to create a triple repeat. As one repeat is two amino acids shorter, variants of all repeats were made with the shorter length. The resulting 64 sequences were aligned and the corresponding phylogenetic tree was calculated. These two were used to generate 16 000 putative ancestral consensus repeat sequences using the FastML server Ashkenazy et al. [2012]. These ancestral sequences were mapped onto the symmetric template using a custom pyRosetta script Voet et al. [2017]. After a short relaxation with the Fast Relax protocol form Rosetta, a ref2015 energy score could be assigned to each sequence. The RMSD from the ideal symmetric backbone was also calculated. Two repeat sequences were selected for experimental validation, the sequence with the lowest energy score (Scone-E) and the sequence with the lowest RMSD (Scone-R).

### 3.2 Purification and Characterisation

The amino acids sequences were reverse translated, optimized for *E. coli* and ordered as synthetic DNA fragments (gBlocks, IDT) afterwards they were cloned into a pET-28b(+) vector and transformed into the BL21 (DE3) strain for protein expression. After induction with IPTG (0.5 mM), the bacteria were grown overnight at 20 °C and pelleted. The proteins were purified following the same protocol as the Cake protein Mylemans et al. [2020a]. Cells were lysed by resuspension in 30 mL buffer 50 mM NaH_2_PO_4_, pH 8, 300 mM NaCl, 10 mM imidazole, 50 ng lysozyme and PMSF (final concentration of 1 mM) and sonication (Branson Sonifier 450, VWR international). Centrifugation was used to separate soluble proteins from cell debris. The supernatant was loaded on a nickel nitrilotriacetic acid (Ni-NTA) resin (Qiagen GmbH) column. The proteins were eluted with a buffer at 300 mM imidazole concentration. After the His6-tag was cut with thrombin, the proteins were reapplied on the Ni-NTA column to remove any remaining impurities. The flow-through was concentrated and loaded on on a HiLoad 16/600 Superdex™ 200 prep grade column (GE Health care) equilibrated with 20 mM HEPES, pH 8, 300 mM NaCl. Protein purity was verified using SDS-PAGE. The fractions corresponding to the main peaks were collected.

1 mg/mL samples of each protein were loaded onto a Superdex™ 200 increase 10/300 (GE Health care) equilibrated with 20 mM HEPES, pH 8, 300 mM NaCl to obtain analytical data.

Circular dichroism (CD) spectroscopy was performed on a JASCO J-1500 spectrometer (JASCO). Protein samples (0.05 mg/mL, in 20 mM phosphate, pH 7.6) were analysed using a 2 mm path quartz cuvette at 20° C. The molar ellipticity was measured from 200 nm to 260 nm. Thermal unfolding experiments were carried out whereby the CD signal at 218 nm was measured as the temperature was increased in steps of 0.2°C

### 3.3 Crystallization and Solving the Structures

Protein samples were concentrated to ca. 10 mg/mL and dialysed overnight against 20 mM HEPES, pH 8 to remove sodium chloride. Crystal screening was performed with a Gryphon robot (Art Robbins instruments, Sunnyvale, USA) using sitting-drop 96-well plates and Screening Suites (Qiagen). Optimization experiments were set up in 24-well hanging drop crystallization plates. The proteins were cocrystallized with silicotungstic acid (STA) by adding equal volumes of proteins at 10 mg/mL and 3.5 mM STA to the screening solution, the compounds of each crystal solution can be found in Table S2.

Crystals were cryoprotected using the crystallization liquor with added glycerol and flash frozen in liquid nitrogen. X-ray diffraction data were obtained at beamline I-04 of Diamond Light Source (Oxfordshire, United Kingdom). Diffraction images were processed with Dials Winter et al. [2018], and scaled with AIMLESS Evans and Murshudov [2013], part of the CCP4 suite Winn et al. [2011]. Molecular replacement with Phaser was attempted using the designed structures, but this failed. Afterwards molecular replacement was tried with smaller fragments and the Cake9 (PDB entry 6tjh) and Cake8 (PDB entry 6tjg) proteins. Replacement with Cake8 and two repeat fragments was successful. Refinement was performed with phenix.refine Adams et al. [2010] and by manual model building with Coot Emsley et al. [2010]. An anomalous map could be created from the diffraction measured at 1 Å. The restraints for the lacunary Keggin POM are based on the ligand structure of the parent silicotungstic acid Keggin (3-letter identifier = SIW). The structure and restraints were edited using REEL Moriarty et al. [2017], removing one W-O moiety to form the lacuna, and the resulting CIF was used for structure refinement to describe the lacunary silicotungstic Keggin (3-letter identifier = LKE). Diffraction and refinement statistics can be found in Table 1.

**Table 1:** Data collection and refinement statistics of X-ray structures. Values in parentheses are for the outer shell.

|  | Scone-R | Scone-E | Scone-E + STA a | Scone-E + STA b |
| --- | --- | --- | --- | --- |
| **PDB entry** | **7awy** | **7awz** | **7aw0** | **7aw2** |
| **Data collection** |  |  |  |  |
| Diffraction source | I04, DLS | I04, DLS | I04, DLS | I04, DLS |
| Wavelength (Å) | 0.9786 | 0.9786 | 0.9786 | 0.9795 |
| Resolution range (Å) | 46.56 - 1.50 (1.53 - 1.50) | 43.2 - 1.60 (1.63 - 1.60) | 71.17 - 2.20 (2.27 - 2.20) | 94.41 - 2.10 (2.15 - 2.10) |
| Space group | $P2_1$ | $P2_1$ | $P6_122$ | $I4_122$ |
| $a, b, c$ (Å) | 40.08, 79.83, 47.23 | 46.44, 61.95, 54.06 | 88.27, 88.27, 194.99 | 133.5, 133.5, 70.44 |
| $\alpha, \beta, \gamma$ (°) | 90, 100.98, 90 | 90, 111.53, 90 | 90, 90, 120 | 90, 90, 90 |
| Reflections (measured/unique) | 212, 215/46, 828 | 166, 347/37, 687 | 601, 413/43, 141 | 421, 666/37, 562 |
| Completeness (%) | 99.8 (99.6) | 99.9 (99.9) | 100.0 (100) | 100 (100) |
| Mean $I/\sigma(I)$ | 16.8 (2.4) | 14.1 (2.6) | 16.7 (3.0) | 11.1 (4.9) |
| Multiplicity | 4.5 (4.6) | 4.4 (4.1) | 12.7 (11.3) | 22.5 (23.4) |
| $R_{pim}$ | 0.019 (0.278) | 0.025 (0.244) | 0.039 (0.53) | 0.077 (0.437) |
| CC(1/2) | 0.999 (0.864) | 0.999 (0.866) | 0.999 (0.919) | 0.951 (0.964) |
| Wilson B factor (Å<sup>2</sup>) | 20.16 | 20.08 | 42.16 | 18.94 |
| **Refinement Statistics** |  |  |  |  |
| $R$ factor/$R_{free}$ | 0.1625/0.1908 | 0.1785/0.2146 | 0.2076/0.2398 | 0.1782/0.2060 |
| No. of atoms in structure |  |  |  |  |
| Protein | 2, 477 | 2, 512 | 2, 449 | 2, 305 |
| Ligand | 18 | 8 | 63 | 105 |
| Water | 343 | 222 | 119 | 79 |
| R.m.s. deviations from ideal |  |  |  |  |
| Bond lengths (Å) | 0.01 | 0.01 | 0.01 | 0.01 |
| Bond angles (°) | 1.2 | 1.03 | 1.29 | 1.13 |
| Chiral volumes (Å<sup>3</sup>) | 0.08 | 0.07 | 0.07 | 0.06 |
| Ramachandran plot, residues in (%) |  |  |  |  |
| Most favorable region | 98.38 | 97.43 | 95.53 | 97.39 |
| Additional allowed region | 1.62 | 1.93 | 4.15 | 2.28 |
| Average $B$ factor (Å<sup>2</sup>) | 25.2 | 24.4 | 42.39 | 24.94 |

## 4 Results and Discussion

Our design method is based on ancestral sequence reconstruction and a symmetrical template, and was previously used to create the Pizza, Tako and Cake proteins Voet et al. [2017]. A schematic overview is given in Fig. 1. We started from the same four blades of the natural template (PDB entry 3hxj) that were used for the Cake protein design Mylemans et al. [2020a]. Following the modified strategy used to design Ika8, models with three repeats were used to construct a three-fold symmetric protein, instead of a single repeat with nine-fold symmetry. To generate the ancestral sequences two options were considered. Single repeat ancestral sequences could be created from an alignment of the four selected blades and these could be mixed in all combinations to create triple length repeats. However, this would generate over ten billion sequences which would be too computationally expensive. Instead, the order of these four repeats was shuffled and a phylogentic tree was created from the resulting 64 combinations. This tree was used to generate ancestral sequences which could then be mapped onto the symmetric template. After a short relaxation with the Rosetta relax protocol, each template was scored with the ref2015 energy function and the RMS deviation from the symmetrical template was calculated. The sequences with the best energy score (Scone-E) or lowest RMSD (Scone-R) were chosen for experimental validation. Each polypeptide carries three identical repeats of a motif, which itself consists of three similar sequences (about 40 residues long) in tandem. The sequences (presented in Table S1) both show roughly 75% sequence identity to the starting template model (PDB entry 3hxj), and 90% identity to each other.

**Figure 1:**
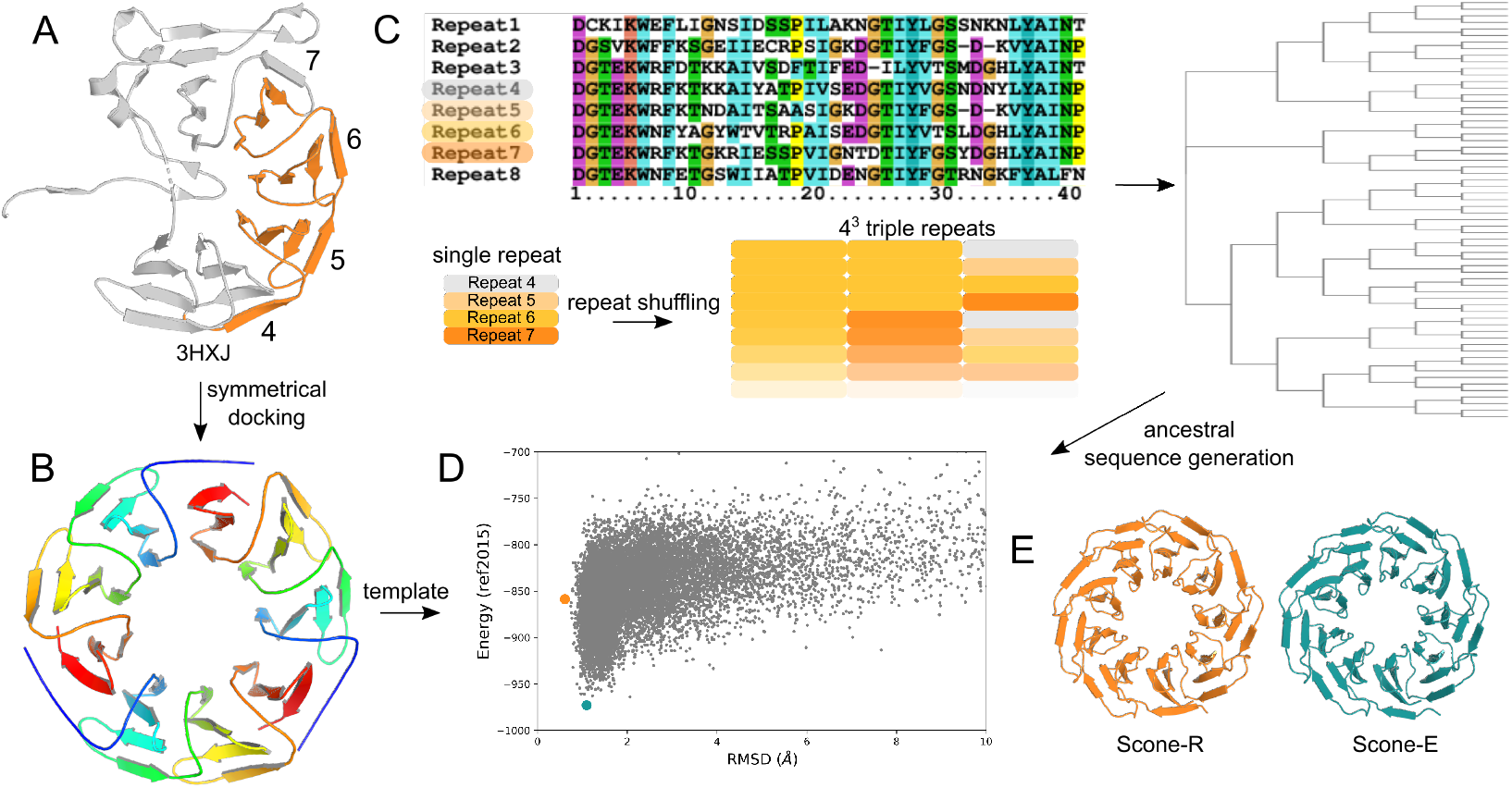
Overview of the computational design method. The (A) incomplete propeller (PDB entry 3hxj) was chosen as template. (B) The triple repeat fragment in orange was extracted and a three-fold symmetric template was created using the SymDock protocol. The repeats were aligned and the sequences of the correctly folded ones were shuffled in all possible combinations. (C) The phylogentic tree of the corresponding 64 sequences was used to generate 16000 ancestral sequences. (D) These sequences were mapped onto the symmetric template with a custom PyRosetta Script. The Energy score and RMSD with the symmetric template were calculated and plotted in function of each other. The lowest energy (teal) and lowest RMSD (orange) sequence were chosen for experimental validation.

Both designs expressed well, and high purity was achieved in two steps using nickel affinity and gel filtration columns. Analytical size exclusion chromatography indicated that the hydrodynamic volumes of both proteins are similar, but smaller than Cake8 or Cake9 (Fig. 2). The CD spectra of Scone-E and Scone-R are identical to that of Cake9, indicating folded proteins with similar secondary structure. Thermal unfolding measurements showed that both proteins are less stable than Cake9, with a melting temperature between 70°C and 80°C (Fig. 2).

**Figure 2:**
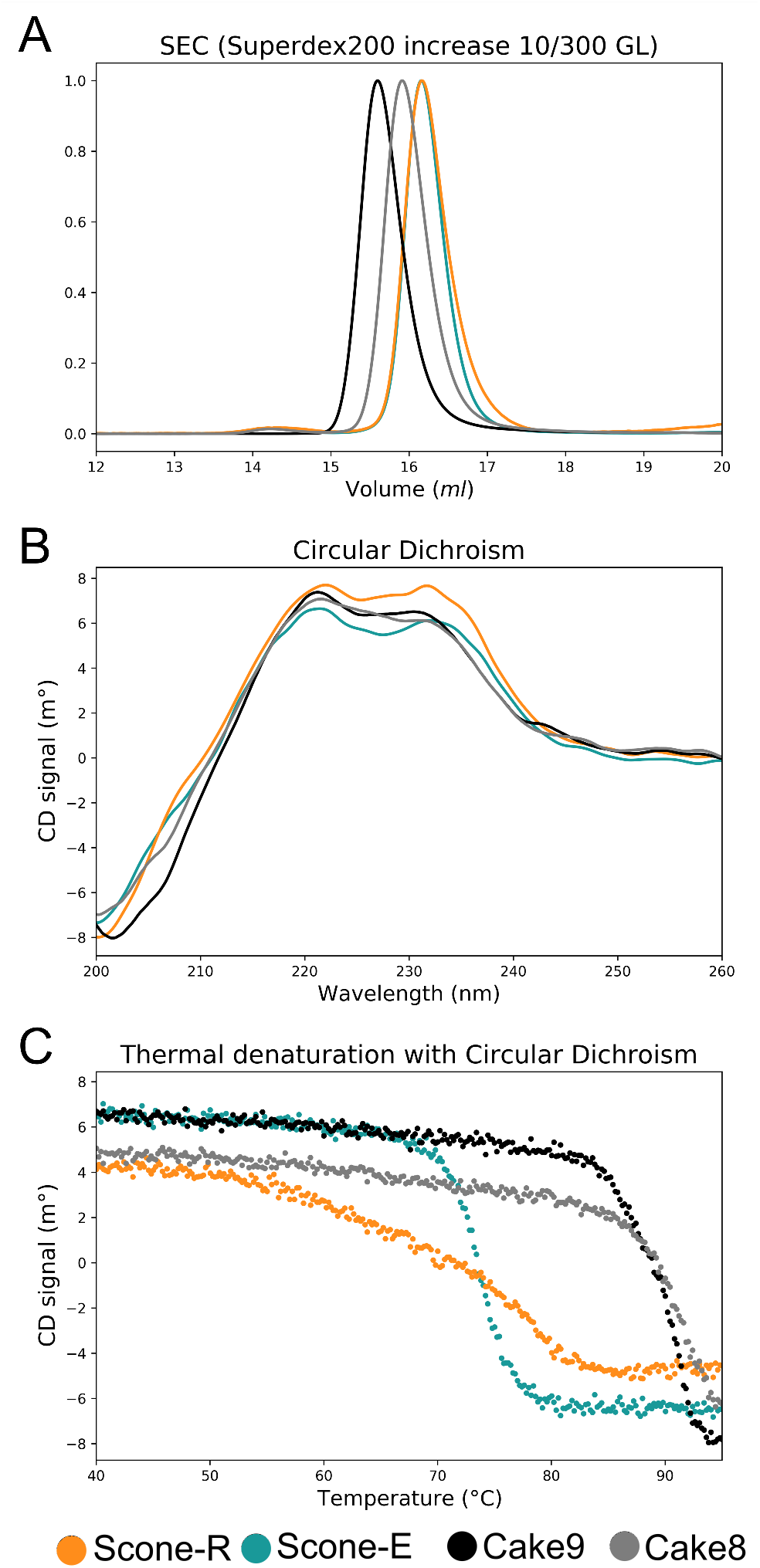
Characterisation of the designed proteins compared to the Cake protein. (A) Size exclusion chromatography shows that both proteins have an identical hydrodynamic volume. They appear to be more compact than both Cake8 and Cake9 although their molecular weight is similar to Cake9. (B) The circular dichroism spectrum of the designed proteins is identical to that of both Cake variants. indicating a similar fold. (C) The CD signal at 218 nm wavelength was measured while increasing the temperature. This showed both proteins are less stable than the Cake variants.

### 4.1 Crystal structure analysis

Scone-R readily crystallized in space group *P*2_1_, allowing data to be collected to a resolution of 1.5 Å, but Scone-E did not yield crystals under any conditions tested in screening. Scone-E has a highly positively charged central cavity that could lead to molecular repulsion, so we attempted to co-crystallize Scone-E with a negatively-charged polyoxometalate cluster (POM). POMs have previously been observed to assist the crystallization of various proteins. Anderson-Evans species are the most used Bijelic and Rompel [2017], Aengus et al. [2018], but phosphotungstic acid Keggin has also been used as a co-crystallization and phasing agent Almo et al. [2007], Ren et al. [2017]. We chose the Keggintype silicotungstic acid (STA) as its symmetry matches the three-fold symmetry the central channel, and it is more stable than other Keggin POMs at physiological pH Zhu et al. [2003], Bajuk-Bogdanović et al. [2015]. Addition of STA to Scone-E yielded three different crystal forms, all of which diffracted x-rays, to resolution limits between 1.6 Aand 2.2 A. Crystals in space-group *P*2_1_ gave the highest resolution data, but were distinctly non-isomorphous with the Scone-R crystals (see Fig. S1). Molecular replacement was initially attempted with the designed models, but this was unsuccessful. Different Cake variants were then tested as search models, and the eight-bladed Cake8 protein proved similar enough to give solutions for both Scone-E and Scone-R. Every crystal form was found to have a single copy of the protein in the asymmetric unit, and refinement proceeded smoothly in each case (parameters are given in Table 1). The models show no remarkable geometrical deviations, with three Ramachandran outliers in Scone-E. Ser34 and Ser152 were found to have slightly unusual backbone geometry in every model. These residues are found at loop regions between strands, with fewer bonds to neighbouring residues and more flexibility. Scone-E was found bound to STA in two crystal forms (called a and b). No POM was found in the *P*2_1_, crystal form. Scone-R and Scone-E overlay closely, with C*α* rmsd between 0.92 and 1.23 Å over 309 ordered residues. The lowest value was found with the crystal form without STA, but the distortions due to the POM molecules are small and localised. The final electron density map unexpectedly revealed that instead of the nine-bladed design, both proteins adopt an eight-bladed architecture with the remainder of the chain invisible in the density. A side-by-side comparison of the designs and experimental crystal structures is shown in Fig. 3. In the majority of *β*-propeller proteins a 1-3 “Velcro” closure is observed in which one N-terminal strand on the outside complements the inner three C-terminal strands of one blade. This configuration is present in Cake, and Scone was designed to share the same feature. In the crystal structure, however, this is not observed, and the ends of the ordered part of the protein chain are found in different blades (4).While the “non-Velcro” configuration does occur in natural propeller proteins, it is rare and has only been reported for the family of prolyl-oligopeptidases Rea and Fülöp [2006]. The absence of strand exchange between blades is even more surprising in the light of recent studies using circularly-permuted Pizza6 protein, which showed this arrangement to be the least stable Mylemans et al. [2020b].

**Figure 3:**
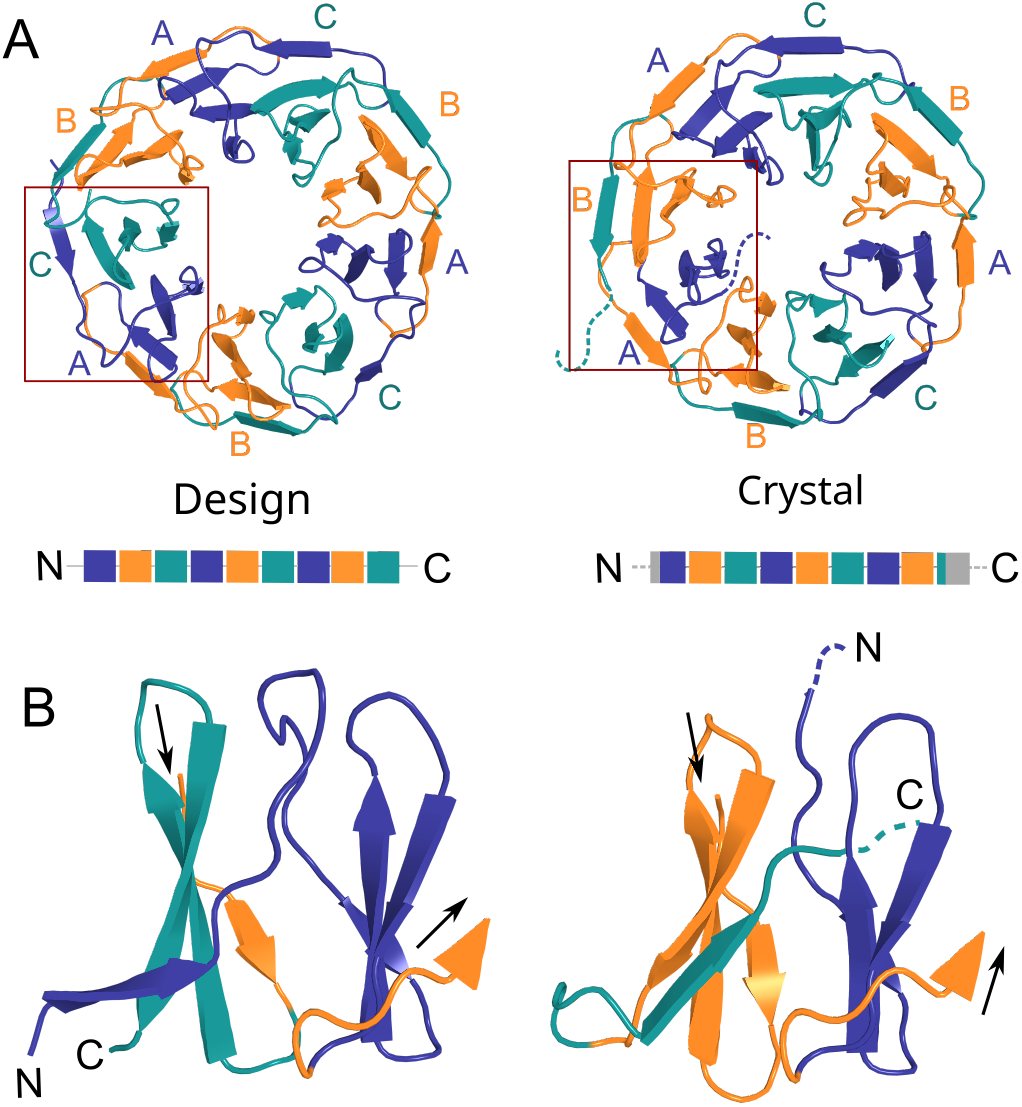
Comparison between design and crystal structure. (A) The designed model is shown with each repeat coloured individually, repeats with the same sequence share their colour, A purple, B orange and C teal. On the right side the crystal structure is shown following the same colouring by repeat. (B) A Close-up of the two terminal blades shows that the protein clearly folds as an eight-bladed propeller instead of a nine-fold one, with parts of both termini not visible in the electron density shown as dotted lines.

Scone-R and Scone-E were intended to form nine-bladed propellers with three-fold symmetry repeating a three-bladed ABC motif (see Fig. S2 and S3). The same C-A interface was expected to form between the first and last blades as between the C and A blades adjacent in the linear protein sequence. Both proteins however formed an eight-bladed propeller missing the final C blade, creating a new B-A interface between the first and last (8th) blade of the propeller. We submitted the sequences to modern protein structure prediction servers. Both I-Tasser Yang et al. [2015] and EVcouplings Hopf et al. [2018] predicted an eight-fold propeller but they also did so for the sequence of Cake9. We excluded the structures of all Cake proteins and the template protein (PDB entry 3hxj) for these predictions to avoid prior knowledge influences. This suggests they are biased towards the much more common eight-bladed fold and are not yet able to distinguish between eight- and nine-bladed propellers. AI based algorithms such as Alphafold Senior et al. [2020] might give an improved prediction.

Using the Rosetta interface energy protocol Bazzoli et al. [2017] and Ref2015, the same scoring function used for the design, we calculated the energy of each inter-blade interface in the expected models and crystal structures (see Table 2). In the design models, the energy of the C-A interface is noticeably higher than those of the A-B and B-C interfaces, giving an uneven energy distribution around the propeller ring (see Fig. S4). In the crystal structures this is not observed. The missing blade changes the overall arrangement of the blades within the propeller, slightly rearranging the interaction between all blades and leading to comparable interactions with very similar interface energies. The proteins therefore adopt a pseudo-symmetric eight-fold symmetrical structure that is more rounded (when viewed along the symmetry axis) than the intended propeller, which had a slightly triangular shape. Hints of strain are apparent in the design models. Average residue energy scores were calculated to be −3.0 Rosetta Energy Units (REU) for the designed nine-bladed symmetric models versus the more favourable −3.5 for the eight-bladed pseudosymmetrical structures. In comparison, the successfully designed Cake9 and Cake8 both have a score of −3.7. Clearly the designed nine-bladed propeller model was not the lowest energy configuration available to the protein. In practice, one ninth of the polypeptide remained unfolded and did not contribute any stabilising energy to the structure. One explanation is that the blades of the template protein (PDB entry 3hxj) inherently favour an eight-bladed organisation, therefore the amino acids necessary to enforce the nine-fold symmetry by stabilising the transition interface between the C and A motif, may not be present in the sample of ancestral sequences. Although the structure of Cake9, which was also derived from the same template, suggests any such bias is unlikely to be very strong. This failure of the design is probably due to the backbone template, which formed a notably distorted propeller with a triangular appearance. This resulted in unfavourable subunit interfaces of the trimeric design, leading to a very uneven distribution of energy between the blades, as well as a high energy overall.

**Table 2:** Internal interface energies between blades in the design and the crystal. Letters for blades follow same convention as seen in Fig. 3

| | Inteface A-B ($REU$) | Interface B-C ($REU$) | Interface C-A ($REU$) | Interface B-A ($REU$) |
| --- | --- | --- | --- | --- |
| Scone-E design | $-47 \pm 4$ | $-58 \pm 3$ | $-39 \pm 3$ | / |
| Scone-E crystal | $-48 \pm 7$ | $-54 \pm 9$ | $-50 \pm 3$ | -45 |
| Scone-R design | $-44.1 \pm 1.8$ | $-55 \pm 8$ | $-34 \pm 2$ | / |
| Scone-R crystal | $-32 \pm 20$ | $-53.3 \pm 1.8$ | $-55 \pm 3$ | -42 |

These bad energy interfaces might be caused by larger deviations from the average blade orientation. This suggests that the propeller blades might have evolved to conform within limits to the average, allowing them to fit between specific neighbours and maintain an overall average curvature which might be one of the reasons why only single blades could be identified as repeat units by a sequence bioinformatics study of *β*-propellers Chaudhuri et al. [2008].

### 4.2 Polyoxometalate interactions

Scone-E could only be crystallized in the presence of STA, and the polyoxometalate could be located in the electron density maps of forms a and b. The tungsten ions were identified by strong peaks in the anomalous scattering map (see Fig. S5). The additive is found at three different positions on the protein surface, and therefore unlikely to bind tightly to any one site, contrary to the strong interactions observed between Pizza6-S and STA Vandebroek et al. [2020]. Instead it is bound to the unstructured loops in crystal form a (grown at pH 7.5), stabilizing a fragment of the final disordered repeat. In all other crystal structure these termini are too disordered and invisible in the electron density. In crystal form b (grown at pH 6.0), two STA molecules are bound to highly positively-charged patches on the protein surface, one inside the protein cavity and one between neighbouring propellers, close to a two-fold crystallographic axis, and therefore modelled at half-occupancy. Both crystal structures with STA bound to the protein are in space groups with higher symmetry than the structures without, due to the crystal contacts formed by the POM (Fig. 5).

**Figure 4:**
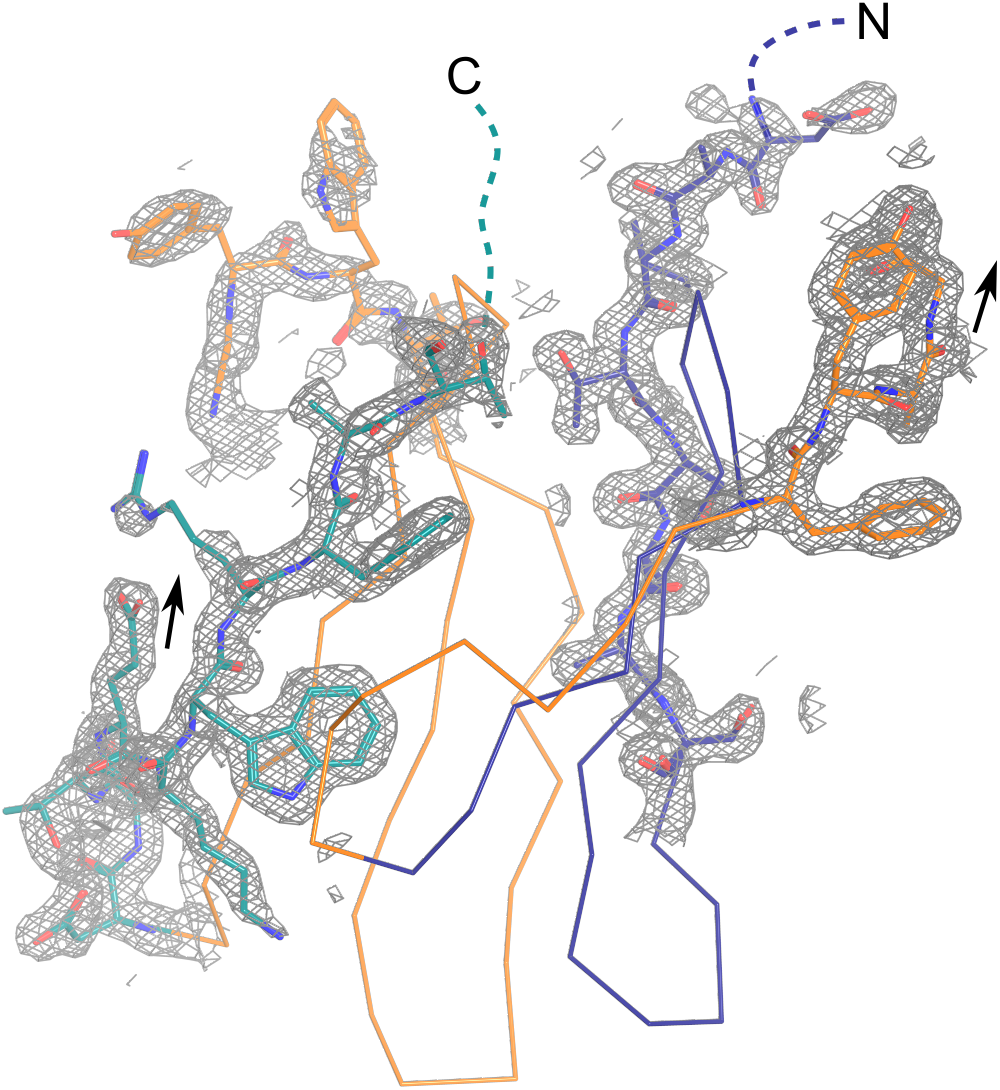
The interface between the N-terminal blade and C-terminal blade is shown. The 2Fo-Fc electron density map at sigma 1.5 is visualized as a mesh clearly showing that the final residues on either termini are unstructured. Arrows indicate the direction of the protein chain from N- to C-terminus

**Figure 5:**
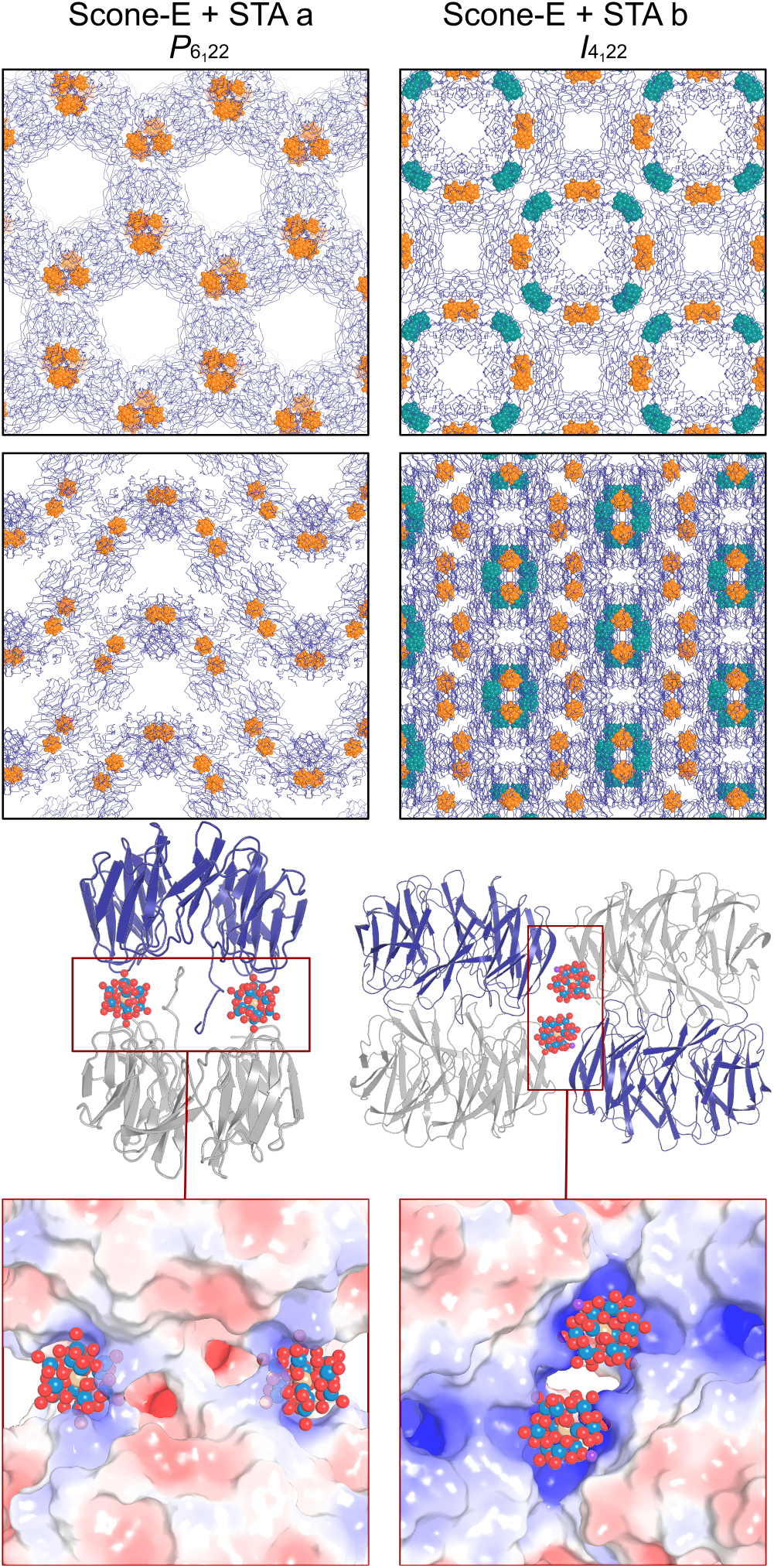
The crystal packing and the role of STA in providing an interface between the Scone-E proteins. When STA is bound in the crystal, the symmetry increases. Below a close-up of the area around the STA is shown with sticks of residues within 4 Å. The whole protein is visible as electrostatic potential surface calculated with APBS at the correct pH. In crystal form a, the POM interacts mostly with arginines in the tail end of the unstructured part of the chain, this allows the protein to form a dimer. In crystal form b, two POMs are present. The STA inside the cavity is attracted by the arginines and lysines in the cavity. This might neutralise the charge of the protein facilitating the formation of a dimer. The POM on the side connects with three protein chains on a positively charged cluster of arginines, lysines and histidines, which might lower the threshold for the proteins to combine as a tetramer

The symmetrical nature of POMs and ability to bind positively-charged surface side-chains on proteins allows them to facilitate crystallization. While STA was required to grow the monoclinic crystal form of Scone-E, it is not found in the final structure, despite the enormous scattering power of the metal atoms within it. Apparently STA can help bring the protein molecules together without becoming bound to them itself. Interestingly, while in previous crystal structure reports STA was observed in its non-lacunary state, here the POM in form b is observed as a monolacunary species with a sodium substituting the tungsten, see Fig. 6. This is not surprising as the Keggin species are known to be pH sensitive. FT-IR measurements have shown that the parent silicotungstic acid Keggin is highly stable at acidic pH, from pH 1 to 6, but starts to show signs of decomposition in aqueous environments starting from pH 6.4 Bajuk-Bogdanović et al. [2015]. It is to be expected that the local environment of the protein, with a localized pKa, can further influence the pH and either protect from or promote POM decomposition. This gives rise to the observation of different POM species at different locations on the protein surface in the same crystal structure.

**Figure 6:**
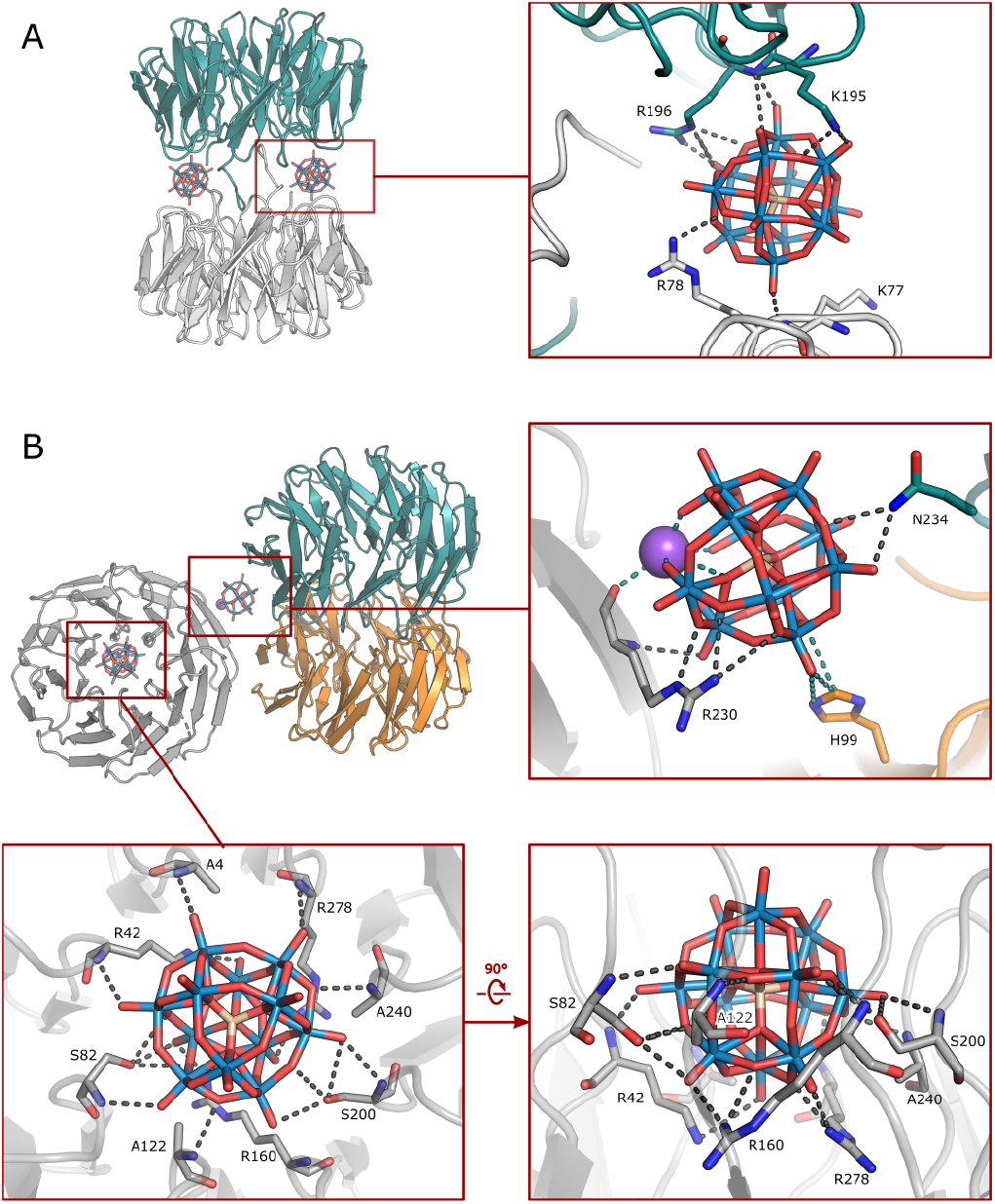
Detailed view of the interactions between the POMs and Scone-E proteins for crystal form a (A) and crystal form b (B). The Scone-E proteins are shown as cartoons, where the light grey chain corresponds to the asymmetric unit; the teal and orange proteins are symmetry equivalent molecules. The interacting residues are shown as sticks, with H-bonding interactions shown as dark grey dashes and electrostatic interactions shown as teal dashes. The POM molecules are shown as sticks, where oxygen is red; tungsten is blue; and silicon is wheat-coloured. The potassium ion bound to the lacunary POM and Scone-E is shown as a purple sphere.

## 5 Conclusions

In conclusion, the two novel designed proteins presented here, Scone-E and Scone-R, adopt a different backbone architecture from the one intended. Like octarellin Figueroa et al. [2016], which was designed as a TIM-barrel but produced a Rossmann-like fold, the Scone proteins are experimentally found to produce stable, soluble structures amenable to crystallisation. They represent interesting cases where in practice the expected model is not the lowest energy configuration available to the polypeptide, and which may help to improve computational design procedures. Interestingly the majority of recent *de novo* designed complexes are very rich in *α*-helices, yet failed designs are rich in *β*-strands. As a guideline for further designs we propose to use a symmetric template with the same symmetry as the number of blades and only vary the sequence as was done for the design of Ika8. Alternatively three-fold symmetric template might be used with a different algorithms not dependent on sequence information such as cost network optimization algorithms Simoncini et al. [2015]. Furthermore, addition of the Keggin polyoxometalate, STA was shown to facilitate crystal growth. Previously the Anderson-Evans species has been used as a crystal adjuvant Aengus et al. [2018] because it is a more pH stable POM. Yet even though the STA became lacunary, it still assisted in crystallization. As POMs are highly interesting for biomedical research yet are not bio-compatible, Bijelic et al. [2019] the Scone-E with the molecule bound in the cavity may form an interesting platform for the further design of bio-compatible POM hybrid proteins as the STA is largely buried inside the protein. This could be achieved by further introducing positively charged and/or hydrogen bonding residues to create a specific high affinity complex.

## 6 Acknowledgements

We would like to thank Els Deridder for helping with the cloning of the constructs. We thank the beamline scientists at the Diamond Light Source macromolecular beamlines for their kind assistance. JRHT thanks OpenEye Scientific Software for financial support. ARDV thanks Research Foundation Flanders for financial support (G0E4717N, G0F9316N and G051917N). BM thanks Research Foundation Flanders for a fellowship (GBM-D3229-ASP/17), LV thanks KU Leuven for PDM fellowship.

---

**Table S1:**
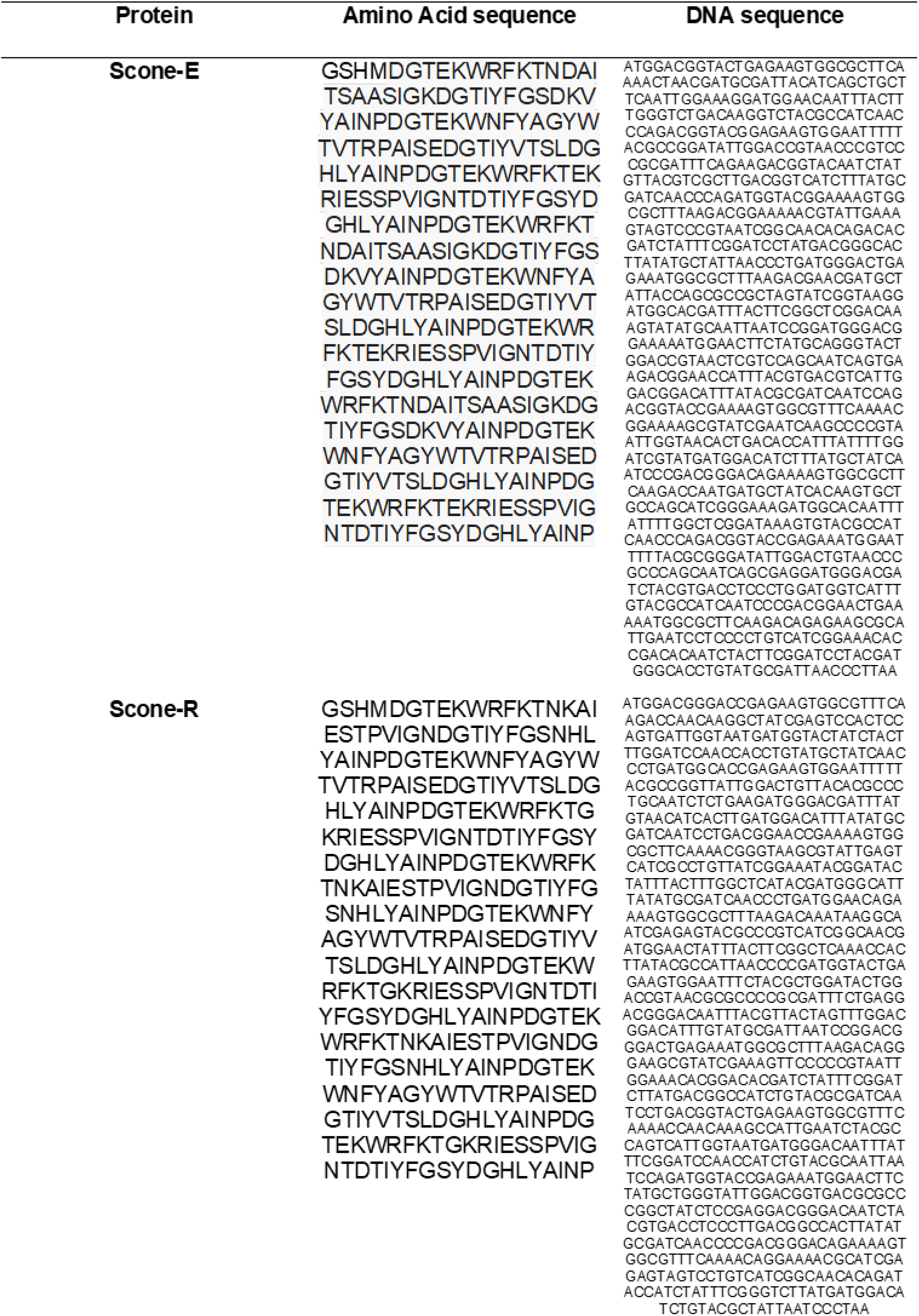
Amino acid and DNA sequences

**Table S2:** Crystallization conditions

| Name | Protein<br>Concentration<br>(mg/ml) | STA concentration<br>(mM) | Reservoir solution |
| --- | --- | --- | --- |
| Scone-R | 10 | 0 | 0.1M Magnesium Acetate<br>0.1M Sodium Acetate<br>4% (w/v) PEG8000 |
| Scone-E | 10 | 3.5 | 0.1M MES pH 6.0<br>40% (v/v) MDP |
| Scone-E + STA a | 10 | 3.5 | 1.8M Na/K phosphate, pH 7.5 |
| Scone-E + STA b | 10 | 3.5 | 1.9M Sodium malonate, pH 6.0 |

**Figure S1:**
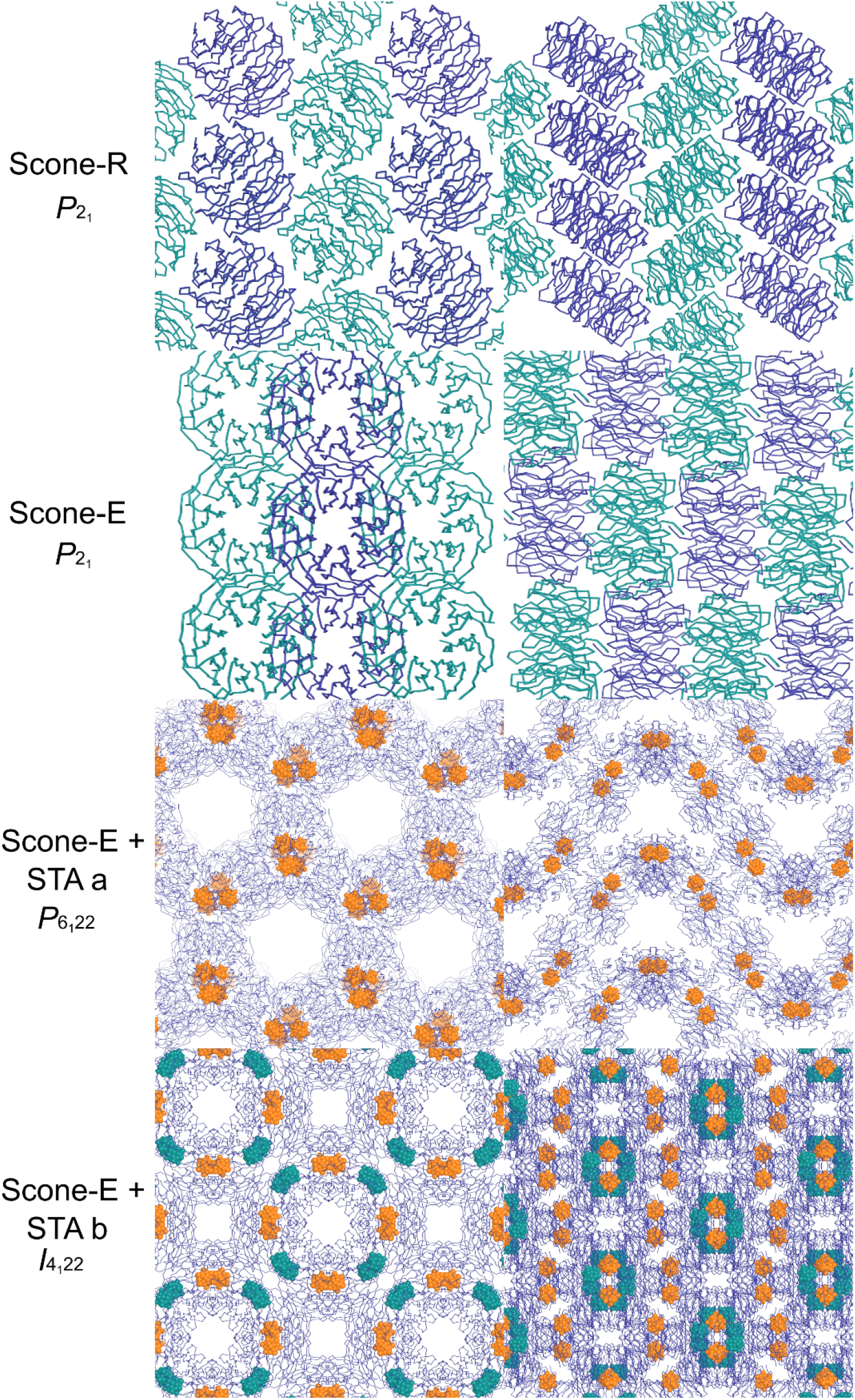
The crystals without bound POM have the same space-group but are not isomorphous

**Figure S2:**
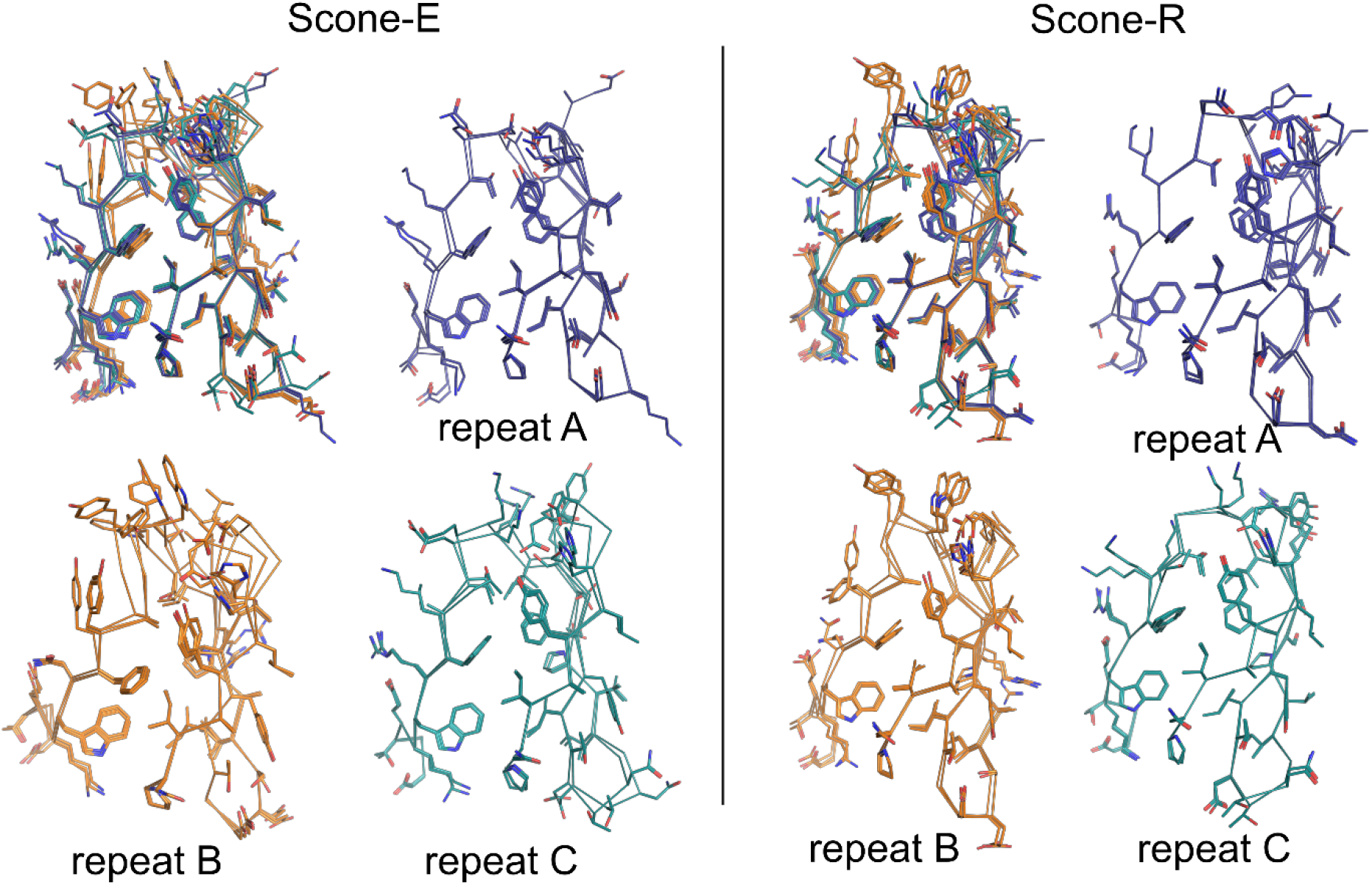
Structural Alignment of repeats: Each repeat is aligned, all together an with the identical sequence repeats separately.

**Figure S3:**
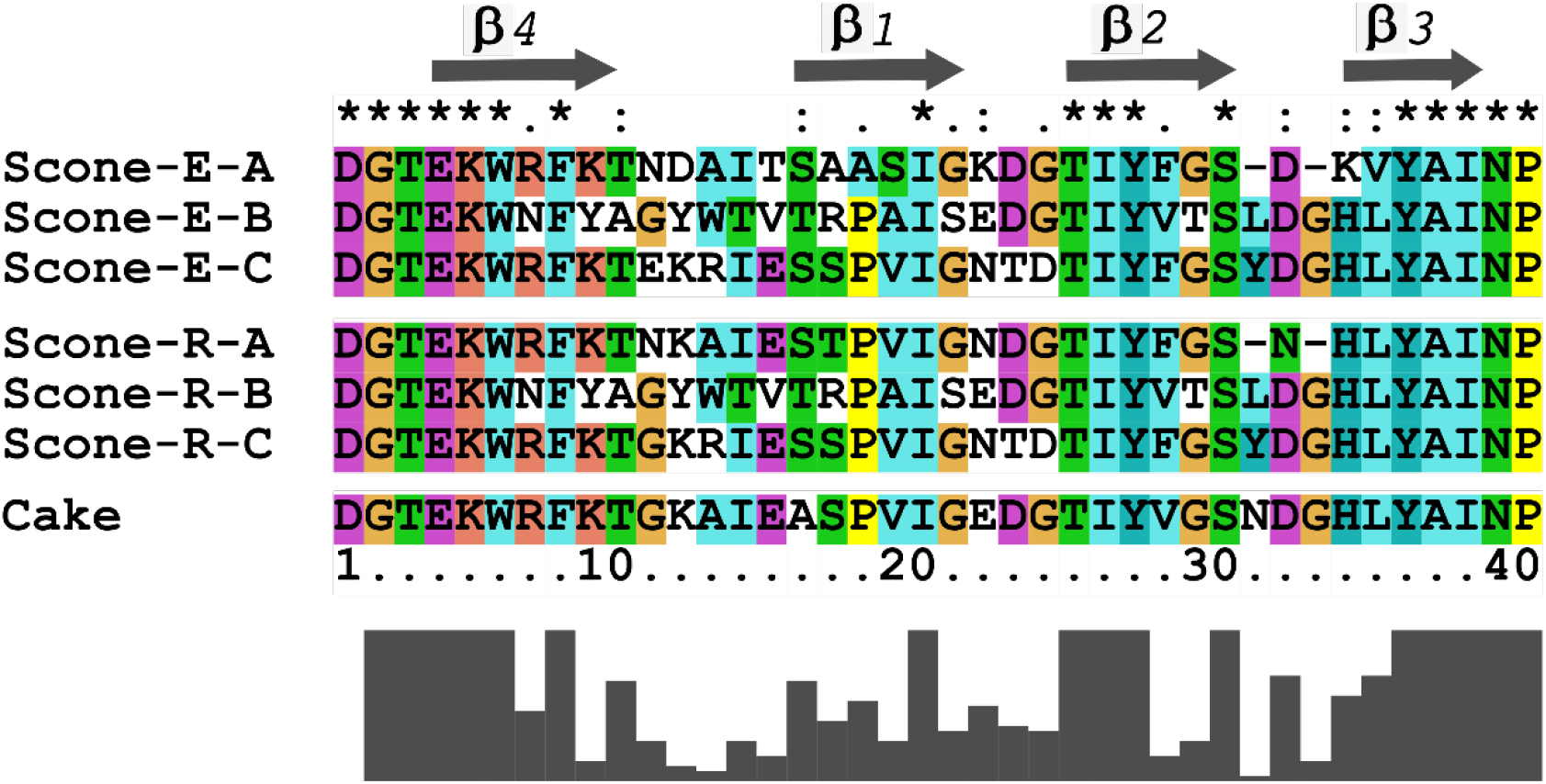
Sequence alignment of repeats: The sequences of the designed Cake protein (PDB:6tjh) is included. On top the location of *β*-strands in the repeat is shown.

**Figure S4:**
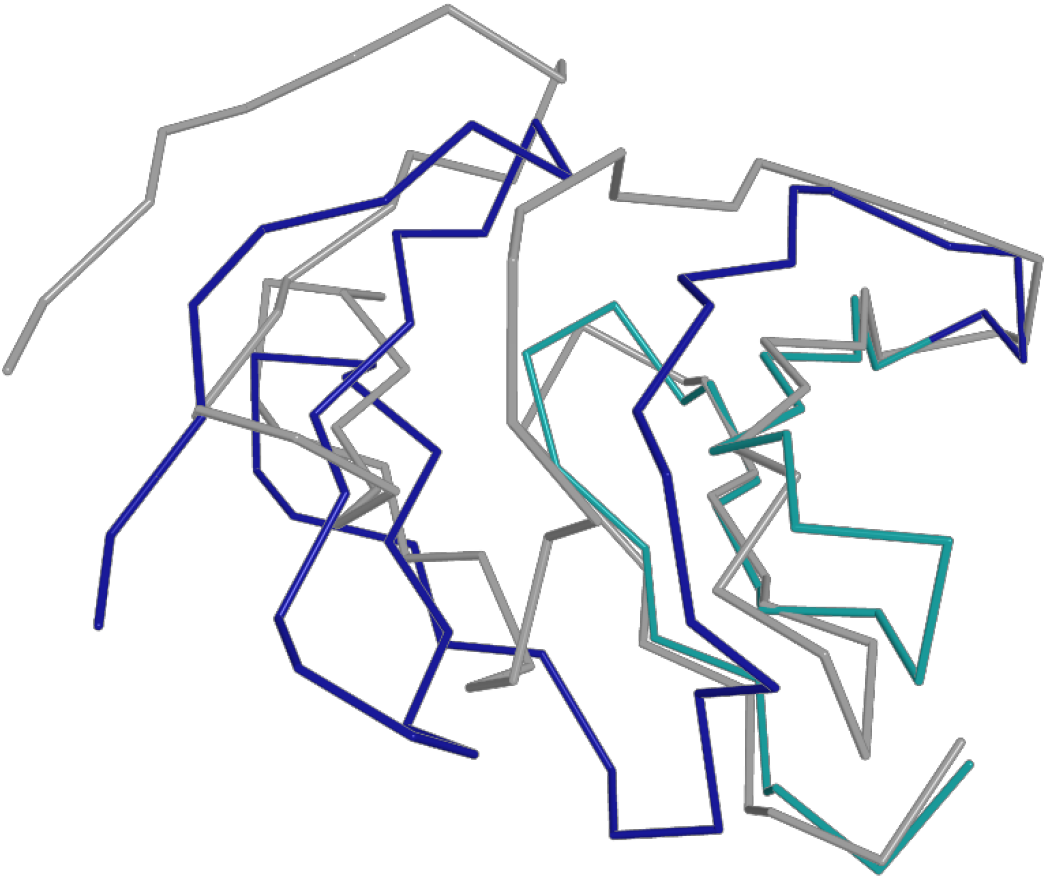
C-A interface: An alignment between the designed C-A interface (grey) and the crystal C (teal) - A (blue) interface shows the clear difference caused by the different fold. The C-blade of the design is aligned on the C blade of the crystal structure.

**Figure S5:**
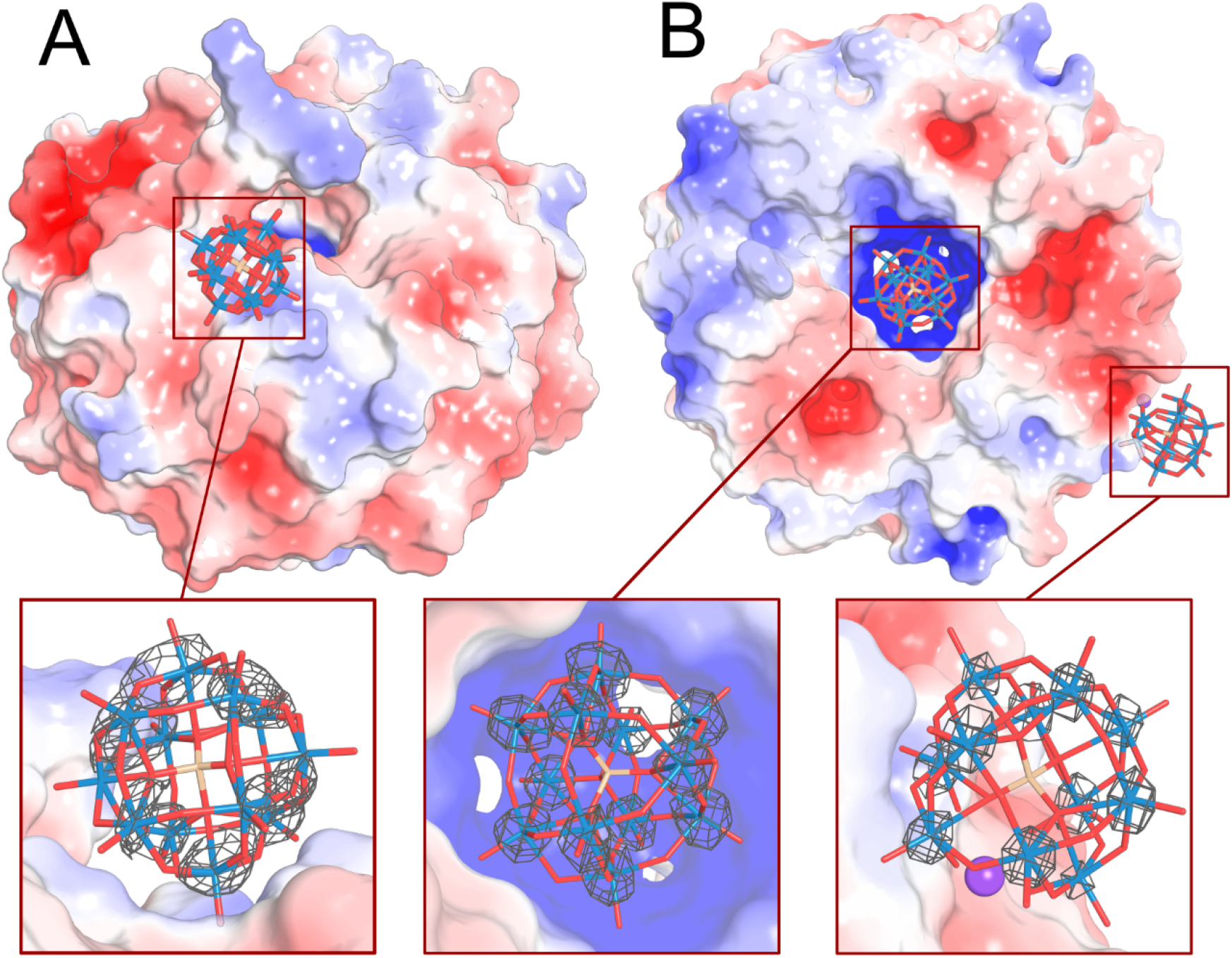
Electrostatic potential map of Scone-E with STA: The electrostatic potential map was calculated with the APBS plugin in PyMOL. Zoom-in on the STA binding site is shown. The anomalous map at sigma 5.5 clearly shows the location of the tungsten atoms.

## Notes

### Competing Interest Statement

The authors have declared no competing interest.

